# Neuronal Population Models Reveal Specific Linear Conductance Controllers Sufficient to Rescue Preclinical Disease Phenotypes

**DOI:** 10.1101/2020.06.01.128033

**Authors:** Sushmita L. Allam, Timothy H. Rumbell, Tuan Hoang Trong, Jaimit Parikh, James R. Kozloski

## Abstract

**Objective:** During the preclinical phase of drug development, potential drug candidates are often screened for their ability to alleviate certain *in vitro* electrophysiological features among neurons. This ability is assessed by measuring treatment outcomes using the population mean, both across different cells and different animals. The go/no-go decision for progressing a drug to a clinical trial is then based on ‘average effects’, yet these measures may not be sufficient to mitigate clinical end point risk. Population-based modeling is widely used to represent the intrinsic variability of electrophysiological features among healthy, disease and drug treated neuronal phenotypes. We pursued a method for optimizing therapeutic target design by identifying a single coherent set of ion channel targets for recovery of the healthy (Wild type) cellular phenotype simultaneously across multiple measures. Specifically, we aimed to determine the set of target modulations that best recover a heterogeneous Huntington’s disease (HD) population of model neurons into a multivariate region of phenotypic measurements corresponding to the healthy excitability profile of a heterogenous Wild type (WT) population of model neurons.

**Methods:** Our approach combines mechanistic simulations with populations modeling of striatal neurons using evolutionary algorithms for population optimization to design ‘virtual drugs’. We introduce efficacy metrics to score population of model outcomes and use these to rank our virtual candidates.

**Results:** We found that virtual drugs identified using heuristic approaches performed better than single target modulators and those derived from standard classification methods. We compare a real drug to the virtual candidates and demonstrate a novel *in silico* triaging method.

## Introduction

The high attrition rate of central nervous system (CNS) drugs is often attributed to off-target activity of lead candidates on neuronal ion channels leading to safety and efficacy concerns. Patch-clamp electrophysiology provides a direct way to measure biophysical properties of ion channel activity and effect on neuronal function. Pharmaceutical companies apply electrophysiology to characterize new leads’ effects on neuronal ion channel activity and neuronal and network function, and *in vitro* electrophysiological assays are being implemented for pharmaceutical safety and efficacy profiling (1–3). Electrophysiological assays for screening new cardiac drugs against myocyte ion channel activities and features are routinely performed because altered cardiac rhythms are readily observed, predictable, and life threatening (4). It is paramount in CNS drug development to similarly advance preclinical testing strategies of new chemical entities to prevent adverse drug reactions and address efficacy (5).

It is widely acknowledged that successfully estimating the beneficial or harmful outcomes of treatments in clinical trials is complex and that failures are often attributed to estimating the average effects of the treatment across the population means while not accounting for population heterogeneity (6). In the CNS, the need exists to design effective drugs for individual tissues, within which physiological variability at the cellular level is pervasive (7,8). This need has been addressed previously through statistical tests of differences among healthy, diseased, and drugged phenotypes of neuronal populations pooled across individual neurons and across different animals (9). Given that a single neural tissue can integrate nonlinearly the dysfunction of a relatively small neuronal cohort (11) the urgency to optimize a drugs’ total efficacy for a tissue population is acute. To that end, we employ rigorous theoretical frameworks and mathematical modeling techniques (12–16) and describe here a computational pipeline for discovering ionic conductance changes at the single neuron level, which together increase efficacy penetrance among target responses of a diverse neuronal population to virtual drugs. Our approach represents a method for pharmacological design that simultaneously addresses the heterogeneity of neuronal responses within a tissue while offering a path towards personalizing neurotherapeutics.

Huntington’s disease (HD) is an autosomal dominant genetic disorder caused by an expanded trinucleotide CAG repeat in exon-1 of the huntingtin gene. Phenotypic changes at the single neuron electrophysiological level within the striatum, a deep structure within the basal ganglia responsible for motor co-ordination and cognition, are believed to underly symptomatic changes in motor and cognitive function during HD manifestation (17). Identifying a single therapeutic mode of recovery for a dysfunctional neural tissue also undergoing neurodegeneration, such as striatum in HD, is challenging, owing to the biophysical diversity of the single neurons comprising the tissue, their adaptive drivers, electrophysiological set points (18) and the confounding effects cell loss can have on normal circuit function (19). Commonly referred to as medium spiny neurons (MSNs), the principal neurons of the striatum exhibit varied active and passive properties in healthy and diseased phenotypes, exhibit manifestation of electrophysiological phenotypes (20) and are the most vulnerable to insult in Huntington’s disease. Here we present an example of a striatal neuron model population for Huntington’s disease (HD) and for their Wild type (WT) background population (20).

Prior experimental studies have explored alterations to physiological and morphological measures among MSNs. Altered active and passive membrane properties such as resting membrane potential, rheobase, input resistance, and firing rate were quantified across D1 and D2 cell types, different HD animal models and healthy and HD phenotypes (21–23). MSNs have been central to studies of pharmacotherapies, such as inhibition of phosphodiesterases (PDEs) of the CAMP and CGMP pathways, aimed at alleviating the above membrane properties both *in vivo* and *in vitro* (9,24). How these pathways engage ion channel proteins via DARPP-32 substrate modulation (25) and regulate the membrane activity in WT and HD phenotypes has yet to be fully understood. Prior *in vitro* and immunofluorescence studies in HD transgenic mouse models detected decreased K+ channel proteins (Kir2.1, Kir2.3, Kv2.1) in MSNs (26). However, it remains unclear if these membrane protein changes are sufficient to explain the altered electrophysiological properties and subsequent vulnerability of MSNs in HD.

Here we present a generic modeling framework that enables building populations of models (PoMs) with characteristics of healthy and diseased neuronal phenotypic categories. We demonstrate parallel approaches to combine statistical and machine learning methods and uniquely identify virtual drugs, which target ion channels and rescue the disease population phenotype towards a healthy phenotype. We also show how scoring these virtual drugs based on their ability to rescue the disease phenotype may be performed based on both heterogeneity and divergence of the neuronal PoMs comprising the phenotype.

Current model-based regulatory evaluations have mainly centered around PK/PD simulations, but with growing impetus for inclusion of *in vitro* electrophysiological assays for safety and efficacy screening of CNS drugs (5), our methods complement triaging strategies to optimize therapeutic target design, which remain areas of strong interest among the pharmaceutical industry.

## Methods

### 1.1 Model and simulations

The MSN model used in this study was published in (27) having been derived with modification from a model published previously by a different group (28). The model comprises a single compartment with eight active ionic conductance models and three specific ionic leak current models. The modifications and channel equations are listed and elaborated in the Supplementary Methods, S1.1.

We constructed two separate databases of PoMs representing the WT and HD phenotypes. To construct the PoMs, we used an evolutionary optimization algorithm, described in detail in S1.2. The optimizations accessed and varied 11 parameters: ionic conductances 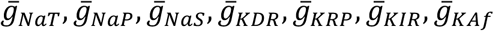 and 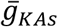 adapted from (28), and GHK based permeability coefficients P_Cl, leak,_ P_Na, leak_ and P_K, leak_. The optimization algorithm (29) was configured to produce models whose features fell within either the WT or HD feature ranges specified (Table 1.1). The parameter conductance’s minimum and maximum ranges were constrained identically for both the WT and HD phenotype-targeted optimizations (Table 1.2).

For the final analysis and virtual drug design, we selected only models that had zero error score (i.e. had feature values within the target ranges (Table 1.1) for all features, which were therefore considered ‘good models’. In addition, we ensured by inspection that every model within the databases of phenotypes exhibited realistic spiking behavior of MSNs by verifying against several stimulation protocols. (see Results).

### 1.2 Virtual drug construction

To identify a coherent target modulation to rescue neuronal excitability in the HD PoM (population of models), we designed virtual drugs (i.e., a set of ion channel parameter perturbations of the HD models) using several statistical and applied machine learning methods:

1. **Single target perturbation method** used the difference between the mean parameter values of transient sodium conductance and fast inactivating A-type potassium conductance of WT and HD PoMs to construct virtual drugs vd_Nat and vd_KAf.
2. **Linear regression method** used the parameter coefficients calculated from regressing the HD PoM’s features against model parameters within a system of linear equations that is solved to compute conductance parameter changes predicted by the regression to transform the mean HD feature values into the mean WT feature values and construct the virtual drug vd_LIN.
3. **Support Vector Machine method** determined the classification boundary between the two phenotypes using a support vector machine (SVM) classifier. SVM is a supervised machine learning algorithm that finds data points closest to a boundary that separates data sets and uses them to help define a linear decision boundary (hyperplane) between data categories. The Python sklearn-SVC package was then used to compute the vector that is normal to the hyperplane separating the two PoMs in parameter space. This vector was then used to effect desired phenotype changes from the HD to the WT PoM and to construct the virtual drug vd_SVM.
4. **n-dimensional histogram method** is a heuristic that took the differences between parameters of each of M members of the WT PoM and parameters of each of N members of the HD PoM’s phenotype to construct M × N difference vectors. Difference vectors were binned in the vector space, according to the size of the differences in each parameter. The mode of this multidimensional histogram was then used to effect desired phenotype changes from the HD to the WT PoM and to construct the virtual drug vd_HIST.
5. **Difference of means method** applied the difference between the parameter value means from **Single target perturbation method** to all 11 model parameters composed a difference vector used to effect desired phenotype changes from the HD to the WT PoM and to construct the virtual drug vd_DIFF.

For each virtual drug, treated HD PoM’s parameters were each modulated uniformly by adding the vectors obtained for each of the above virtual drug construction methods and are termed HD+*virtual drug*.

### 1.3 Scoring Metrics

We used the following metrics to quantitatively compare the performance of each of the above virtual drugs’ effects in rescuing the neuronal excitability of the HD PoM’s features:

1. **Euclidean Distance 3D** measures the distance between centroids of two populations of data using three membrane properties: V_m_, R_m_ and Rheobase.
2. **Jensen-Shannon Divergence** measures similarity between two probability distributions within the same three-dimensional space of membrane properties. Note this metric is a symmetrized version of Kullback-Leibler divergence, and its square root is the Jensen-Shannon Distance. The probability distribution of WT PoM’s features and recovered HD PoM’s were first estimated using the kernel density estimation method (KDE) from the Python scikit-learn package (sklearn-learn). We approximated from WT PoM’s (n=1219) the multinomial distributions using KDE for a maximum of 3 features and used these KDEs, and the Python SciPy package (30) to quantify the Jensen-Shannon Distance and recover the HD PoM’s (n = 1223).
3. **Total Score** measures the mean of normalized Euclidean Distance 3D and Jensen–Shannon Distance metrics. We used this score to quantify the efficacy of different virtual drugs for recovering the WT PoM from the the HD PoM.
4. **Euclidean Distance 7D** measures the distance between the centroids of the WT and HD population data phenotypes derived from seven of the membrane properties listed in Table 1.1 (excluding FR100 and TFS100).
5. **Models Retained** counts the number of models within the HD PoM that satisfied the additional constraint by inspection of retaining realistic spiking behavior of MSNs after virtual drug perturbations.

### 1.4 Other visualization and statistical methods

For analysis of the resulting HD+*virtual drug* PoM, we employed the following:

1. **Convex hull method** used the Python scipy.spatial convex hull package to calculate the hull vertices and edges bounding the 11 data points obtained from Beaumont et al., (9) by means of the quick hull algorithm (http://www.qhull.org). This hull is illustrated in Fig. 1 for the three categories WT, HD and HD+PDE10i.

**Figure 1:**
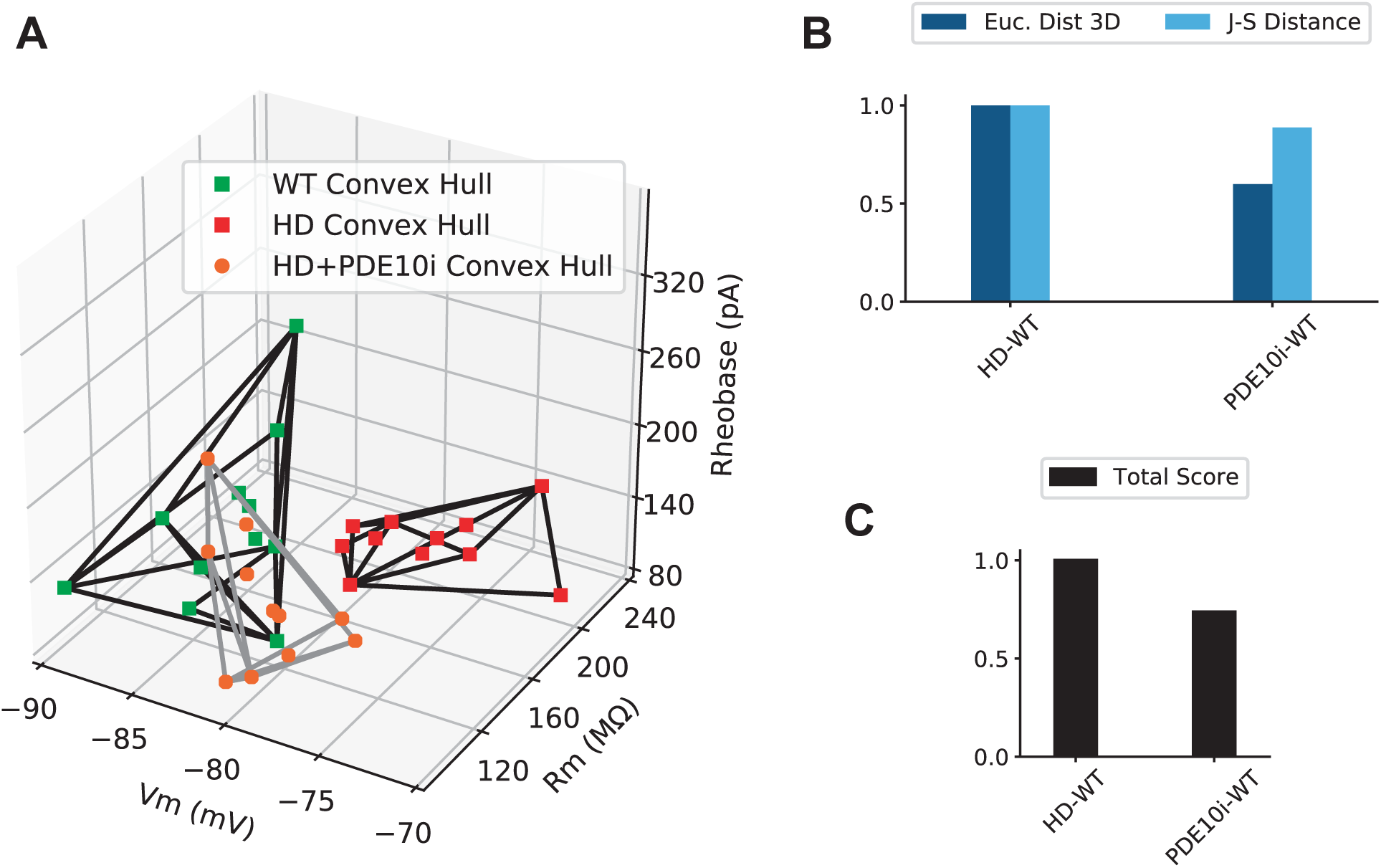
Efficacy evaluation metrics. **A)** Convex hulls enclosing WT, HD and PDE10i treated HD phenotypes in three dimensions representing membrane properties of MSNs illustrate location of each phenotype in feature space. **B)** Distance metrics used to score the performance of the drug for its ability to recover the HD phenotype in feature space (right bars) quantified as a proportion of the original Euclidean distance and J-S Distance between WT and HD PoMs (left bars; normalized to 1.0). **C)** PDE10i treated HD phenotype’s Total Score is the average of Euclidean Distance and J-S Distance.
2. **K-nearest neighbor search** applies a method for finding a predefined number of points closest in distance to the new point based on standard Euclidean distance. We set nearest neighbors to 2 in order to derive at least two ‘matched models’ for every real empirical data point in Figure 2A.

**Figure 2:**
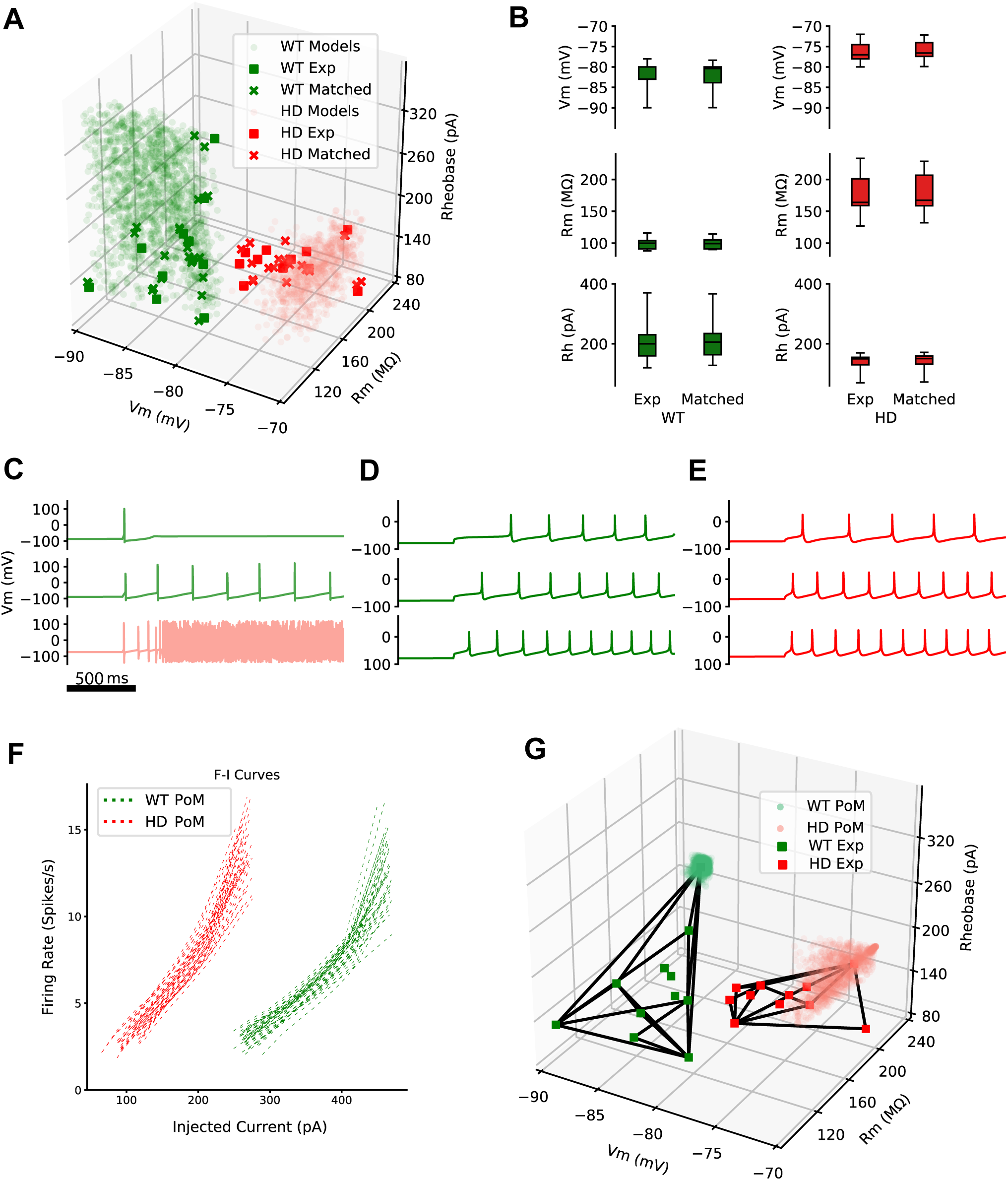
Population of models to characterize WT and HD phenotypes. **A)** Evolutionary algorithm generated PoMs for WT and HD phenotypes. Light circles show PoMs, solid squares show experimental data, and the subset of each PoM matched to the empirical data using k-nearest neighbors search are shown as ‘x’. **B)** Box plots of the three membrane properties of the matched models and empirical data are similar (two sample K-S test statistic for WT V_m_ (p=0.63), R_m_ (p=0.63), and rheobase (p=0.96); HD: V_m_ (p=0.82); R_m_ (p=0.96); Rheobase (p=0.83)). **C)** 50% of matched models exhibit unrealistic spiking behavior. Top and middle example models from the WT PoM shown in green. Bottom example model from the HD PoM shown in red. D) and E) Addition of further spiking feature constraints to ensure the PoMs exhibit realistic spiking patterns of WT **(D)** and HD **(E)** phenotypes. **F)** F-I curves of WT (green) and HD (red) PoMs show heterogeneity within each PoM, and lowered excitability in HD. **G)** Additional spiking constraints collapsed the feature space for the PoM of the WT phenotype, thus setting a narrower target range for reversing the heterogeneous HD PoM’s phenotype.
3. **Statistical methods** to calculate p-values between the empirical and matched models were calculated using the Python scipy.stats package’s Kolmogorov-Smirnov (K-S) statistic applied to two samples.

## Results

Thus far, there have been few drugs approved for treatment of HD, with most treating chorea and movement disorders (31). Some antiepileptic drugs are prescribed for various neurological disorders (including HD) and target neuronal excitability through modulation of ion channels. The main mechanisms of action of these drugs, in addition to targeting voltage-gated ion channels, are modulation of glutamatergic and GABAergic neurotransmission and intracellular signaling pathways (32). Due to the preclinical success of a PDE10 inhibitor (i.e., the PDE10i known as PF-230920) in rescuing *in vivo* and *in vitro* neurological deficits in HD symptomatic animal models, the path to clinical trial was taken for this PDE10i, but the drug failed to ameliorate motor and functional disturbances in HD patients (9,33). Therefore, the failure of the Pfizer Amaryllis trial made clear that finding a drug candidate based on current preclinical criteria triaging methods does not necessarily predict the successful outcome of subsequent human clinical trials. Therefore there exists not only a need for better drug design, but also for better means of preclinical scoring and deciding which drugs should enter clinical trial, such that trials which maximize translatability and ability to mitigate clinical end point risk of failure might become possible (34).

### Metrics for scoring efficacy of phenotypic recovery

We used the pre-clinical *in vitro* data from Beaumont et al., (9), precursors to Pfizer’s Amaryllis trial, in order to quantify using our multivariate methods how well the drug recovered compromised neuronal excitability in the Q175 disease phenotype model. To assess the extent of separation in the space of the measured neuronal phenotypes (single neuron excitability features) we used convex hulls, which provide the smallest convex set of enclosure of points for a data set (35). In Fig. 1A, we show convex hulls that visualize the enclosures of data from the WT (green) and HD (red) phenotypes in a three-dimensional (3D) feature space, comprising passive membrane properties of medium spiny neurons (MSNs). The phenotype of MSNs rescued by the pharmacotherapy of the PDE10i is represented by the convex hull of HD+PDE10i (orange), which intersects the WT convex hull space. We quantified the performance of the drug by extracting our two 3D metrics from the data for the distance between both HD and HD+PDE10i phenotypes and the WT phenotype, Euclidean distance (ED3) and J-S Distance (see methods), and here report values normalized to HD-WT distances for comparison. The resulting HD-WT distances were normalized to 1.0, while the drug treated phenotype HD+PDE10i, the ED3 was calculated as 0.6 and the J-S distance as 0.92 (Fig. 1B). Next, we calculated the mean of these distance metrics to find the ‘total score’, which for HD+PDE10i was 0.73 (Fig. 1C). We conclude that based on these preclinical data, PDE10i was advanced as a viable HD drug candidate based on a recovery of ∼27% of the total score derived from these distance metrics.

### Population of models for characterizing WT and HD electrophysiological phenotypes

We wondered if a computational model of the MSN (27,28) might allow further refinement of our characterization of the preclinical data from Beaumont et al., 2016. Creation of large databases of model neurons can provide insight into how neuronal membrane response properties are determined by the underlying ionic conductances (12) and help elucidate intersubject variability (13). We created a database of WT and HD phenotypes from different instances of the model, thus creating two PoMs. These PoMs reproduced the empirical ranges of membrane properties (Fig. 2). In Fig. 2A, green and red squares represent the original preclinical data from 11 neurons for WT and HD respectively. To create the PoMs, an evolutionary algorithm described in (36) was used (see Supplementary Methods) to explore the model’s eleven dimensional parameter space, with each dimension representing one of eight active ionic conductance and three specific ion permeability parameters (see Table 1.2 for details of parameters ranges used for sampling). Separate optimization runs targeting each phenotype, WT and HD, generated PoMs with output features bounded by specified ranges. The WT PoM comprised 1650 different parameter settings (Fig. 2A, light green circles), while the HD PoM comprised 859 (Fig. 2A, light red circles). Model instances within the WT and HD category spanned the range of all three passive membrane features (Fig. 2A). From this large database of models, we sampled models to generate a joint distribution across all three feature dimensions, comparable to that of the empirical data, using the *k-*nearest neighbor algorithm (see Methods). The sampled models, comprising the 2 nearest models for each empirical observation, are each indicated by ‘×’ among the cloud of all model instances. Box and whisker plots summarize the experimental data from (9), and the sampled models for both WT and HD phenotypes (Fig. 2B) show good agreement between model outputs and the data (two sample K-S tests for WT V_m_: p=0.63, R_m_: p=0.63, and rheobase: p=0.96; HD V_m_: p=0.82, R_m_: p=0.96, and rheobase: p=0.83). We could, therefore, sample model instance feature values from the same distribution as the experimental data. The evolutionary search and subsequent sampling methods allowed us to create PoMs in close proximity to the preclinical data in feature space. This PoM generation framework is robust for creating PoMs representative of different empirically observed phenotypes by varying the parameters of a single underlying mechanistic model.

To examine the spiking properties of the sampled models, we then applied several step current injection protocols. Our optimization had only targeted passive membrane properties, and therefore no specific features of MSN spiking activity, such as AP height, after-hyperpolarization potential (AHP), coefficient of variation of interspike intervals (ISI CV), time to first spike (TFS; also termed ‘spike latency’) and firing rate (FR), were optimized with these properties. Without constraints on active membrane properties, in some cases our optimized and sampled models failed to reproduce spiking features characteristic of MSNs. We observed that 50% of these models entered depolarization block under physiological step current injection. Because the empirical data reported in (9) did not include raw traces from which to extract these spiking features, we decided to make the assumption that the empirically sampled neurons did not enter depolarization block and proceeded to survey the literature to obtain normal measurements for each (21,22), thus establishing their acceptable ranges for our subsequent refining of the MSN PoMs (See Methods, Table 1.1).

A new optimization run targeted the 14 features reported in Table 1.1. This optimization found 1219 WT and 1223 HD models constrained by the complete set of 14 feature ranges. Models had regular firing patterns and none entered depolarization block when injected with current stimuli ranges up to rheobase plus 100 pA (Rh+100pA; Fig. 2D, E). Model outputs each showed characteristic spike latencies that were greater in WT models than in HD models, and and each showed higher firing rates with increasing depolarizing stimulation across both phenotypes as observed in (21,22). Fig. 2F, shows F-I curves obtained from 20 models for each phenotype, constructed using step currents in increments of 20pA from Rh-60 pA to Rh+100 pA. The HD models had a lower rheobase and exhibited hyperexcitability (red in Fig. 2E), whereas the WT models had a higher rheobase (green in Fig. 2E). Heterogeneity of F-I curves are seen across both populations.

The evolutionary algorithm produced PoMs that conformed simultaneously to passive membrane feature ranges of WT and HD empirical observations, as well as generic descriptions of MSN active spiking properties (Fig. 2F). However, these new PoMs failed to capture the full range of empirical diversity among passive properties (V_m_, R_m_, Rh) to the same extent that the original PoM from our prior, less constrained optimization did (Fig. 2A). The new HD PoM (1223 models) was reasonably well spread within the convex hull of HD empirical data (black boundary in Fig. 2F) but, again, not as well as the original WT PoM, and the new WT PoM (1219 models) was clustered at one vertex of the convex hull of WT empirical data and thus did not sample the entire WT convex hull. In general, optimizations of PoMs encounter problems when adhering to too many feature constraints simultaneously. Despite these limits to sampling, we observed that both model populations (WT and HD; Fig. 2F) were significantly different from each other (one-way ANOVA, p< 0.0001). This difference is consistent with the statistical differences among empirical observations of (9). In summary, we constructed a large database of models (∼2400), that obeyed the active properties of MSNs, specifically AP height, AHP, spike latency, firing rates, and ISI CVs. These models also approximated passive membrane properties that distinguished WT and HD MSNs. Furthermore, in subsequent sections of this report, the more narrowly constrained WT models (Fig. 2F) will present an even more challenging target than their empirical counterparts for ameliorating HD models using virtual drugs. We propose that provided the underlying model is a good one, any successful virtual drug identified under these specific, more stringent criteria for resolving HD phenotypic disturbances, should also in general be able to recover the WT phenotype from the HD phenotype.

### Analysis of parameters reveals ionic conductances associated with phenotypic differences

We further analyzed the parameter distributions underlying our PoMs and created a reverse screen for the origins of neuronal excitability differences among the HD and WT phenotypes. Box plots of ionic conductance parameter values comprising the WT (green) and HD (red) phenotype PoMs normalized to the mean of the WT models are shown in Fig. 3A. The most significant difference predicted by the PoMs is a downregulation in KAf conductance, which further corroborates experimental evidence for intrinsic excitability being mediated by Kv channels (37,38). The transient Na conductance was also significantly downregulated in our HD models, and while to our knowledge no direct evidence of reduced expression of Nav1.2 channel proteins in HD exists, there is evidence for sodium channel β4 subunit downregulation in HD transgenic animals, which may underlie neuritic degeneration (39). We calculated a K-S test statistic between WT and HD model parameters and found those parameters for which the K-S distance was ≥0.7: for Nat, 0.95; for KAf, 1.0; for NaP, 0.73; and for KRP, 0.76. The cumulative distribution functions of these model parameters are shown in Supplementary S1. Because relying on mean data alone provides insufficient insights into underlying causes of functional variability (16), we also examined correlations among the parameters by constructing a heatmap representing pairwise correlations between model parameters (Fig. 3B). WT model parameter correlations are shown in the lower triangle of Fig. 3B, and HD model parameter correlations shown in the upper triangle. A strong correlation between Nat and KAf exists (r > 0.8), which is diminished in HD phenotypes (r∼0.3). We next assessed how the ionic conductance parameters were correlated to each of the first 7 features from Table 1.1. We applied linear regression methods (40) (see Supplementary Results) to uncover the relationship between each of the model’s ionic conductances and the features of model output for both phenotypes. The linear regression coefficients indicate the sensitivity of the perturbation of individual conductances and their influence on the relative model properties against which they were regressed. The firing rate property was most sensitive to two conductances: NaS and KRP (Figure S1). These two conductances were highly correlated among both WT and HD models seen in Fig. 3B. Both of these ionic conductances have long time constants of activation, which may underlie their ability to maintain subthreshold excitability and alter repetitive firing rates (41,42). The coefficients that most influenced rheobase, a critical differentiating feature of the phenotypes, were P_KCNK (permeability to K+ ions) in both phenotypes, followed by KAf conductance. This is not surprising, as prior experimental work quantified the effects of potassium permeability on depolarization and intrinsic properties of hippocampal pyramidal neurons by exposing the culture to KCl, which lowered input resistance and resting potential, and increased rheobase (43). Overall, most of the features could be predicted well with linear models (R^2^ values > 0.9; See Supplementary Results) with the exception of spike latency feature for WT, where the goodness of fit measure was low (R^2^ = 0.3), but a polynomial regression fit with order 2 improved the R^2^ value to 0.6 (not shown).

**Figure 3:**
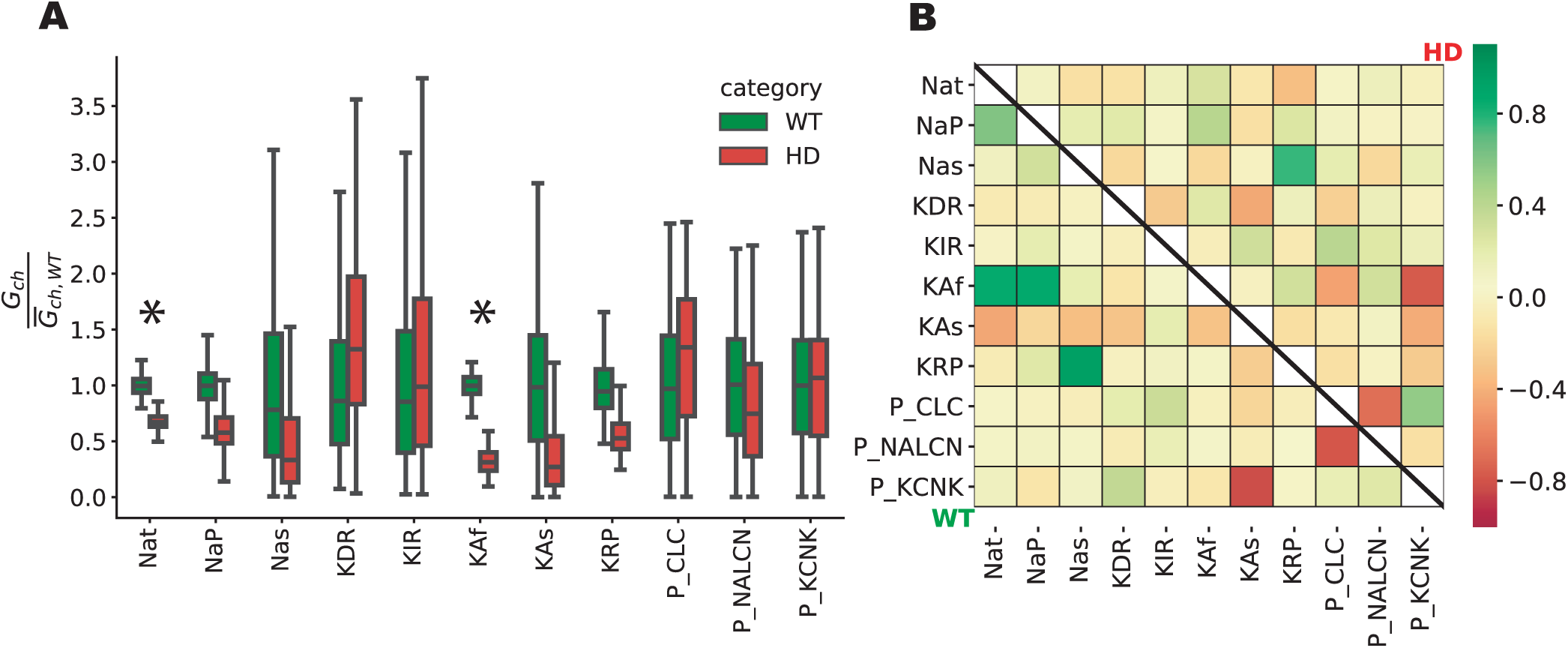
Analysis of parameters of the WT and HD PoMs reveals decreased ionic conductance changes associated with HD pathology. **A)** Box plots of ionic conductance parameter values comprising the WT (green) and HD (red) phenotype PoMs. Values are normalized to the mean parameter values for the WT model. Ionic conductance densities that show significant differences between WT and HD (K-S distance> 0.9) are indicated with “*”. **B)** Correlation matrix of the ionic conductances within the WT PoM (lower triangle) and the HD PoM (upper triangle). Diagonal is left blank. Correlations between certain conductances (e.g., the subthreshold slow sodium, Nas, and persistent potassium, KRP) are conserved across the two phenotypes.

### Virtual Drug Design

While the above described methods that are useful to characterize parameter sensitivity with respect to model features, they are insufficient to reveal a coherent target modulation profile, i.e., a set of perturbations of ionic conductance parameters of the model sufficient to rescue excitability phenotypes in HD. We have termed these unique combinations of ionic conductance perturbations ‘virtual drugs’ and present the various methodologies we evaluated to efficiently recover the HD phenotypes towards the WT phenotype feature space. ‘Efficient recovery’ refers to perturbing the model parameters such that the resulting treated HD models’ output features are in close proximity to the WT models’ features within a multidimensional measurement space, calculated according to our distance metrics. We demonstrate and present the results of these methodologies in the following sections.

### Single target modulation

Fast inactivating potassium conductance, KAf, was significantly downregulated in the HD PoM (Cohen’s d measure > 3). Accordingly, we considered gKAf to be the key parameter, as it was highly correlated to features such as TFS, Rheobase, and FR (see Supplementary Results), and was the strongest coefficient of the first principal component that explained 36% of the variance in the combined WT and HD PoM’s parameter space (not shown). Next in importance was the transient sodium current, Nat, which was critical for regulating AP related features together with KAf (Figure S1). For this reason, our first virtual drugs were constructed by modulating each of these conductances. We applied these virtual drugs to each HD model instance in the HD PoM by changing parameters according to the virtual drug and measured the outcome in the 3D feature space of passive membrane voltage, membrane resistance and rheobase. By way of example, when Nat was modulated, we first calculated the difference between the means of the WT and the HD PoM’s ‘Nat’ conductances, which we used as a reference perturbation in constructing the first virtual drug vd_Nat. The relative magnitude and direction of this reference perturbation is represented in Fig. 4A as the mean conductance of the HD PoM’s parameters after applying (i.e., adding) the reference perturbation to each. We calculated this mean conductance (1.47 for gNat) by dividing the mean of the perturbed HD PoM’s parameters by the mean of the HD PoM’s parameters (virtual drug-modulated parameters are shown relative to the original HD mean parameters). The virtual drug vd_Nat was then applied to each model in the HD PoM at four fractional doses of the full reference perturbation (100%, 75%, 50%, and 25%; Fig. 4C). KAf conductance of the HD PoM were similarly modulated by the second virtual drug vd_KAf (Fig. 4B, D). The resulting phenotypic recovery in 3D feature space in response to vd_Nat at these doses is shown in Fig. 4C, and in response to vd_KAf in Fig. 4D. As shown in this figure, vd_Nat did not recover the HD phenotype, which was not surprising, since Nat was not identified as contributing to variation among the membrane excitability features in the PoM. In contrast, vd_KAf was sufficient to transform the HD PoM towards the WT PoM. We quantified the extent of efficacy and recovery of the HD PoM, summarized using four metrics (See methods). vd_Nat performed more poorly than vd_KAf in recovering the Euclidean distance in 3D, Euclidean distance in 7D, and J-S Distance. In contrast, vd_KAf at 100% dose recovered the Euclidean distance by at least 60% across all features and >75% in the three passive membrane features, but it failed to address the heterogeneity with the HD PoM and resolve its divergence relative to the less diverse WT PoM. Also, vd_KAf applied at higher doses understandably generated excess risk of altering other membrane properties, to the extent of breaking the response of the model, such that resulting features could not be measured. We quantified this effect with an auxilliary measure of the proportion of models retained, which decreased as vd_KAf dose increased, resulting in retention of only 40% of models at the reference dosage. It will be interesting to examine this model-based risk’s relationship to toxicity measures of real CNS drugs in subsequent work.

**Figure 4:**
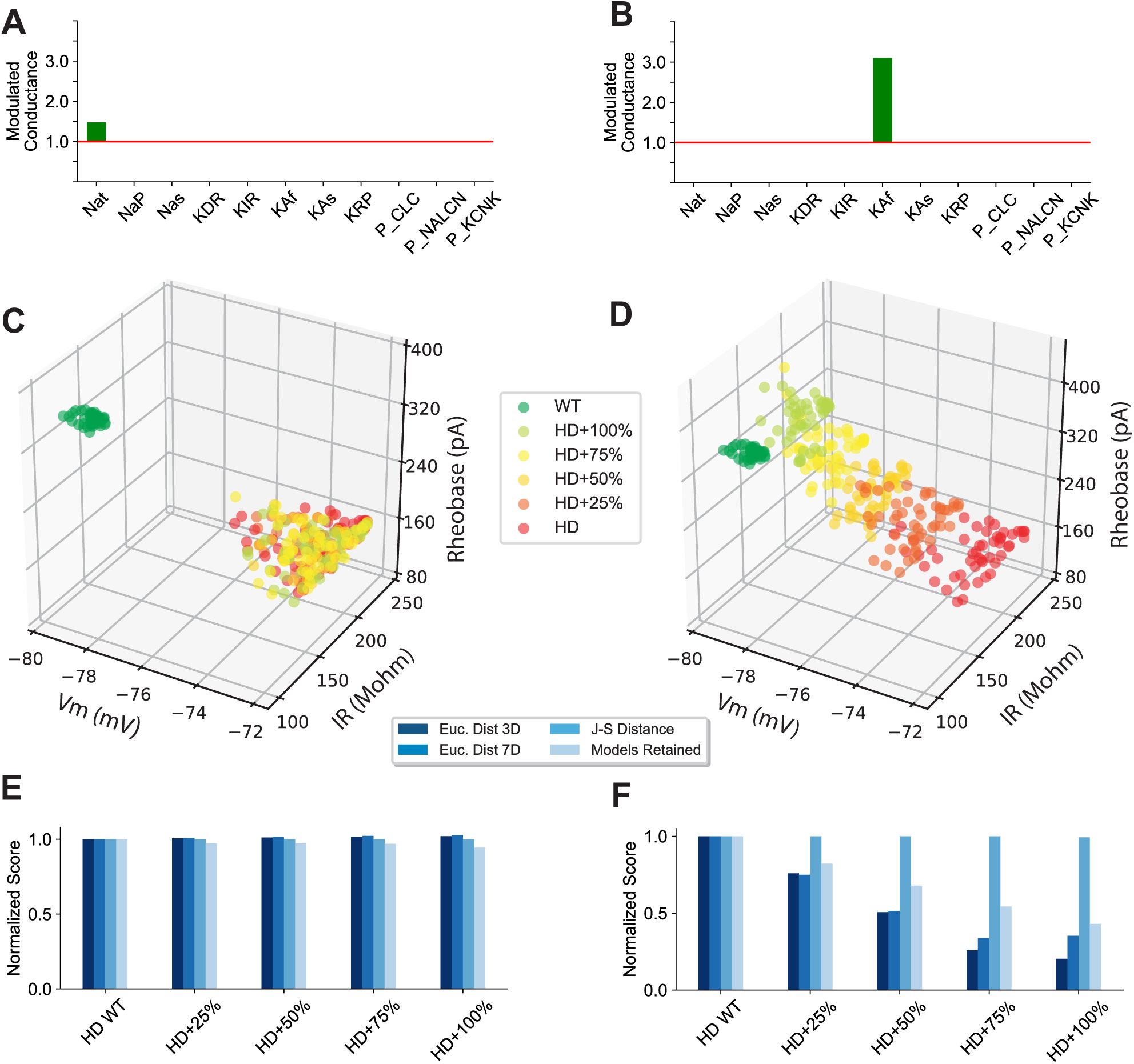
Single target modulation performance metrics. **A)** Single target perturbation to Na^+^ transient conductance (Nat) applied to the HD PoM. **B)** Perturbation of fast inactivating potassium conductance (KAf) applied to the HD PoM. **C)** Phenotypic recovery in three-dimensional feature space with perturbation vector shown in A. **D)** Phenotypic recovery with perturbation vector shown in B. **E)** Four metrics (see color key) score the Nat perturbing virtual drug efficacy for recovering the WT PoM’s phenotypes from the HD PoM’s phenotypes: Euclidean distance in 3 dimensions, Euclidean distance in 7 dimensions, and J-S Distance (see Methods), as well as Models Retained. “Models Retained” refers to models whose features remain within permissible realistic behavior of MSNs after the perturbations are applied at each of the four intermediate doses. **F)** Same as E) but scoring the KAf perturbing virtual drug.

### Multiple target modulation

There has been a shift away from the “one drug, one target” approach, wherein highly potent and specific *single-target* treatments were preferred to polypharmic drugs, as they purported to mitigate off-target side-effects. However, translatability from *in vitro* drug effects to *in vivo* efficacy was poor with this approach (33,44). While single target strategies appeared reasonable for known disorders controlled by single targets, such as in cardiac arrythmias (45), neurological diseases involve disruption of network homeostasis (46) and breakdown of multiple contributing underlying biochemical cascades. For this reason, we aimed to augment polypharmic approaches with our reverse screen and identify highly potent and specific, *multi-target* treatments to alleviate complex phenotypes by the simultaneous modulation of multiple ion channels.

### Linear methods sufficient to recover phenotype

Next, we considered whether it might be possible to modulate multiple parameters simultaneously to more efficiently perturb the HD models towards the WT phenotype and achieve better recovery than accomplished by our single parameter modulations. Linear regression analysis (see also Supplemental Materials) provided insights into how multiple conductances regulated different electrophysiological properties. These linear relationships engaged multiple ionic conductances, supporting the idea that to regulate a specific feature in desired manner, multiple conductance parameters should be perturbed simultaneously. A cartoon representation of how this perturbation is determined for the HD PoM is illustrated in Fig. 5A (See Methods for details on its construction). To explore a second method for constructing a multiple parameter virtual drug, we employed a linear SVM classifier to define a separating axis orthogonal to the hyperplane classification boundary. The projection of the PoMs’ parameter vectors onto the separation axis determined by the SVM classifier then generated a set of values (i.e., ‘scores’ of arbitrary units), histograms of which together with kernel density estimation results are shown in Fig. 5B. These histograms illustrate the clear separation between the two model phenotypes. A similar approach using logistic regression (i.e., the ‘characteristic direction’ method) was used previously to identify similarities and dissimilarities between gene expression studies from multiple experiments, when the underlying parameter space is high dimensional (47).

**Figure 5:**
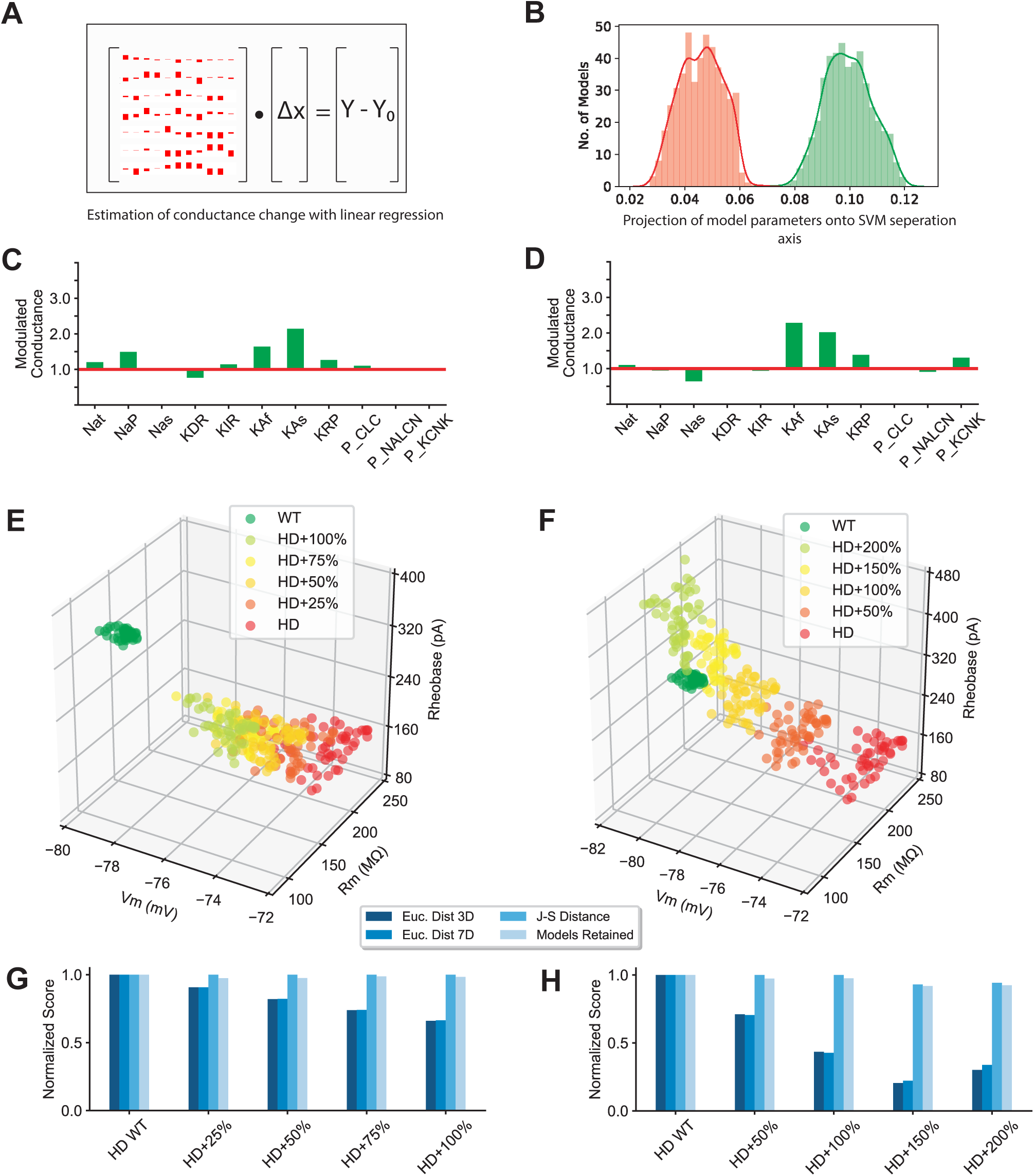
Multiple target modulation performance metrics using linear methods. **A)** Linear Regression method used to determine conductance change for expected feature change (Y-Y_0_). **B)** WT and HD models separated by a hyperplane determined using Linear Support Vector Machine method **C)** and **D)** Modulated conductance by perturbation vectors determined from method described in A and B. **E)** and **F)** Phenotypic recovery in three-dimensional feature space with perturbation vector shown in C and D. Note the difference in dosage for F (see text). **G** and **H)** Four Metrics used for scoring the virtual drug’s perturbation efficacy in terms of its ability to recover the HD PoM in Euclidean distance and Divergence (J-S Distance) when applied with perturbations in C & D at four intermediate doses.

First, we calculated a reference perturbation from the linear regression analysis described in Methods. The conductance vector determined by the regression (the linear solution ‘X’ in Fig. 5A) represents the conductance changes necessary to effect the feature change from the HD to the WT PoM (the right hand side ‘Y-Y_0_’ in Fig. 5A). We used this conductance vector’s magnitude and direction as the reference perturbation for the third virtual drug vd_LIN. The relative perturbation is represented in Fig. 5C as the mean conductances of the perturbed HD after applying (i.e., adding) the reference perturbation to each HD model. The virtual drug vd_LIN was also applied to each model in the HD PoM at additional fractional doses, as above. The resulting phenotypic recovery in 3D feature space in response to vd_LIN at these doses is shown in Fig. 5E.

Next, we calculated a reference perturbation along the separation axis of the SVM analysis described in Methods. This axis is defined by a conductance vector, which is normal to the hyperplane and has a magnitude (1.12) related to the distance between the z-score normalized parameters of the HD and WT PoMs. Specifically, the conductance vector represents the conductance changes necessary to transform the HD PoM’s parameters (in the direction of the WT PoM’s parameters), such that the SVM classifier’s accuracy is reduced by 50%, and to alter the model scores of the HD PoM (x-axis panel 5B) by 0.05 (arbitrary units), as required to classify them as WT PoM. We used this conductance vector’s magnitude and direction for the reference perturbation of the fourth virtual drug vd_SVM. The relative perturbation is represented in Fig. 5D as the mean conductance of the HD PoM’s conductance parameters after applying (i.e., adding) the reference perturbation to each. The virtual drug vd_SVM was then applied to each model in the HD PoM at fractional doses ranging from 50-200%. The resulting phenotypic recovery in 3D feature space in response to vd_SVM at these doses is shown in Fig. 5F.

While vd_LIN at maximum dose only recovered 34% of the Euclidean distance measures of the HD PoM, vd_SVM at the 150% dose recovered 80% of these measures (Fig. 5G, H). The further incremental dose (200%) of vd_SVM moved the PoM away from the WT PoM and worsened the recovery in terms of our distance metrics. As with single target modulation, both approaches failed to reduce the divergence metric, J-S distance, and therefore suffered from an inability to resolve the full cohort of our HD PoM’s members’ dysfunctions, while resolving only the PoM’s mean response metrics.

### Virtual drug with heuristic approaches provided best phenotypic recovery

To address this divergence of our HD PoM’s members undergoing virtual drug treatment from their targeted WT PoM phenotypes, we explored the direct use of the WT PoM’s members to generate statistics on the various directions in parameter space pointing towards each from the HD PoM’s members. We calculated the parametric differences between each HD model and each WT model, which we reasoned could supply the necessary range of modulations needed to yield an effective transformation of the PoM. Our first approach (Fig. 6A) estimated the most frequently occurring range of difference vectors between the two PoMs. We constructed a multidimensional histogram (illustrated for only 2 of those parameters in Fig. 6A) over all parameter differences between members of the WT and HD PoMs (top left). We formulated a second related approach by calculating the differences between the mean WT and HD parameters for each parameter independently, as shown in Figure 6B. While the former method estimates the required perturbation based on every possible parameter difference between all pairs of models across the two PoMs, the latter method estimates a direction in parameter space from the mean values of the WT and HD PoMs’ parameters. Figs. 6C and D, represent the mean conductances achieved by the multiple target virtual drugs vd_HIST and vd_DIFF, applied to the HD PoM. The approaches are similar in their target modulation profile with subtle changes around modification to NaS, KIR and KCNK parameters. Both altered transient sodium and fast inactivating potassium conductances in similar relative proportions, along with NaS and KRP, resulting in feature transformations among the PoM that were remarkably good. Not only did these virtual drugs move the HD PoM to its closest proximity to the WT PoM, but at intermediate dosages, they effected a progressive decrease in the HD PoM’s members divergence from the WT PoM (J-S Distance). This category of virtual drug perturbation was therefore unique in its total efficacy and represents a novel approach to virtual drug design (Figs. 6E, F). Having addressed our most stringent criteria governing efficacy (Figs. 6G, H), these drugs performed surprisingly well and therefore warrant further investigation for designing HD therapeutics.

**Figure 6:**
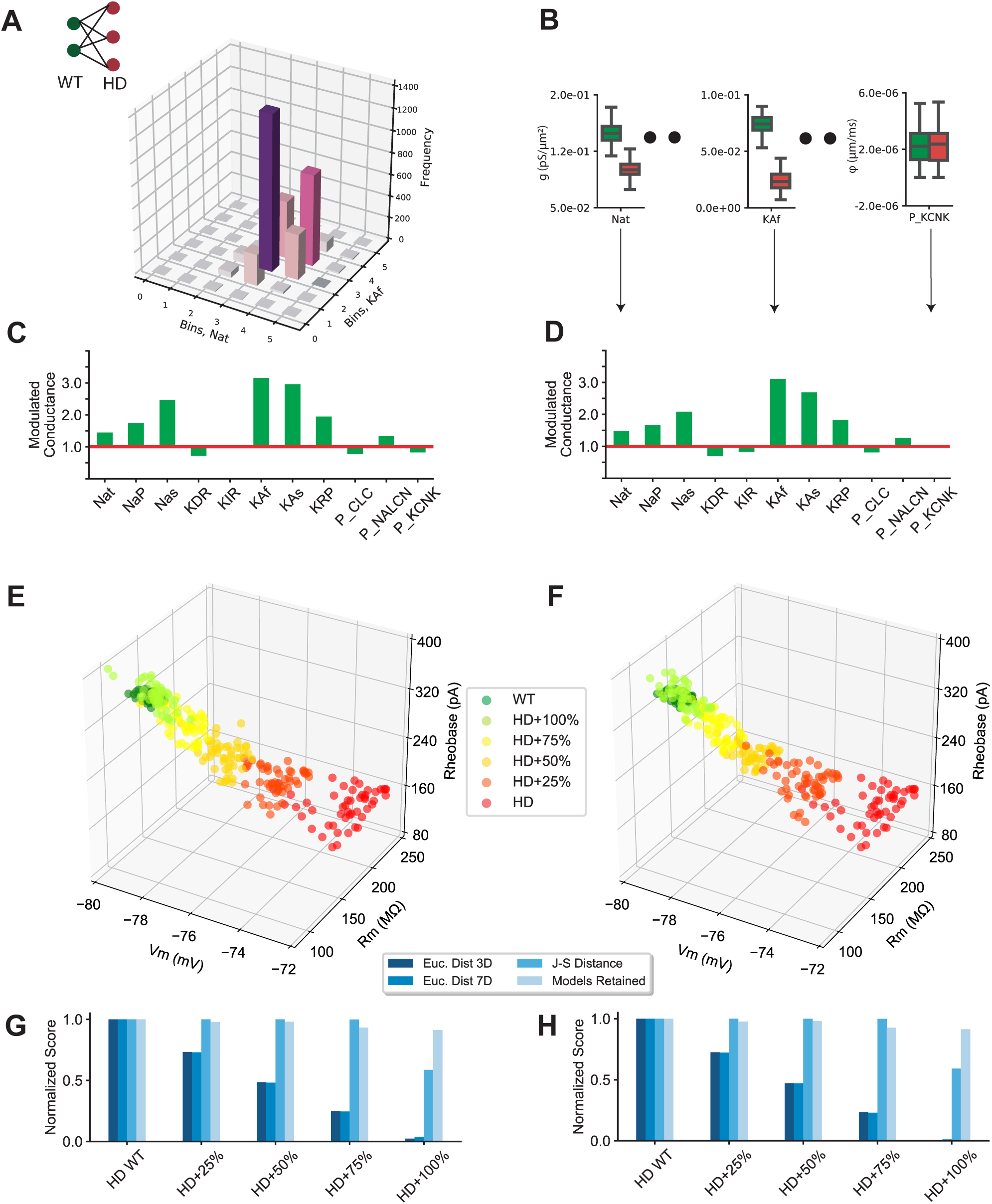
Multiple target modulation performance metrics using heuristic methods. **A)** Multidimensional histogram (shown for only two parameters) of all possible parametric differences between WT and HD PoMs (top left) **B)** Individual Parametric differences between the means of the PoMs. **C)** and **D)** Conductance modulations by perturbation vectors determined from method described in A and B. **E)** and **F)** Phenotypic recovery in three-dimensional feature space with perturbation vector shown in C and D. **G)** and **H)** Four Metrics used for scoring the virtual drug’s perturbation efficacy in terms of its ability to recover the HD PoM in Euclidean distance and Divergence (J-S Distance) when applied with perturbations in C & D at four intermediate doses.

### Triaging approach using virtual drugs to design efficient target modulation

We further compared the performance metrics of all the virtual drugs used in this study. First, we examined the extent of recovery in the distributions of all features. As illustrated in Fig. 7A, feature distributions of the HD PoM (red) appear disjoint from the WT PoM (green), but appear proximal to the WT PoM (green) when perturbed with the best virtual drug vd_DIFF (orange). Fig. 7B provides a visual representation of the rescue of WT phenotype across all feature pairs. As seen in subplots s, t and u for pairwise combinations of V_m_, IR and Rh, the virtual drug vd_DIFF also addressed the divergence in distributions from HD to WT. However, for features such as FR50 and TFS50, the vd_DIFF treated HD PoM distributions exhibited a wider resulting spread. These results allude to possible next generation model enhancements beyond the current single compartment approximation using ionic conductances alone, such as optimizing ion channel parameters governing activation and inactivation kinetics and time constants. Additional structural complexity, such as dendritic compartments, may also be required to address these features.

**Figure 7:**
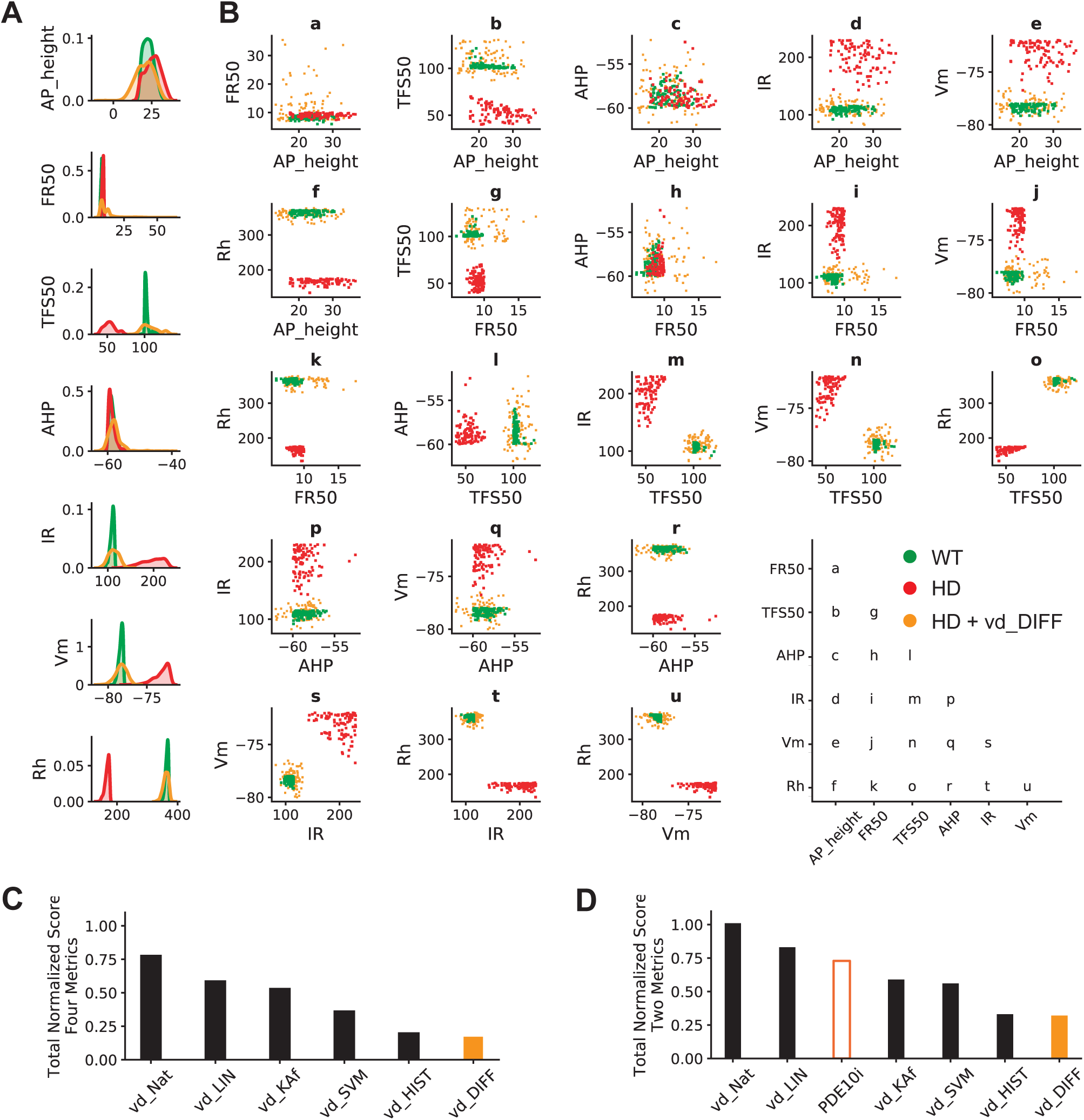
Validation and performance metrics of virtual drugs. **A)** Kernel density estimates show full recovery of the HD PoM (red) when perturbed with the best virtual drug (orange) and overlap the WT PoM (green). **B)** Pairwise scatter plots of all features. The best virtual drug, vd_DIFF, recovered spiking features that were added in this study, and which extended previous empirical measures of excitability. **C)** Drug scores calculated from the averages of four metrics (Figs. 4-6) for each virtual drug at their best dosage identified in this study. **D)** Drug scores calculated from the average of two metrics (Euclidean distance in 3 dimensions, and J-S Distance) allowing comparison across each the virtual drugs at best dosage and the real drug, PDE10i.

We note that virtual screening and scoring methods in traditional drug discovery are mostly centered around estimating ligand-protein binding affinities and energies (48). Scoring drugs based on their ability to address recovery in terms of population heterogeneity and divergence has remained unexplored prior to our study. Here, we have introduced a novel screening method to quantify drugs’ efficacy in terms of recovering not just single models or a population mean, but for their ability to resolve a heterogenous PoM’s multidimensional phenotype, representative of the full contingent of cellular phenotypes occurring in the target tissue. We scored virtual drugs from the average of four metrics (See Methods and Figs. 4-6). The virtual drug vd_DIFF performed best, followed closely by vd_HIST, each of which recovered ∼70% of the total WT-HD distance. We also compared these drugs in two metrics, Euclidean and J-S Distance, for three passive membrane properties (Fig. 1) and found similarly high efficacy.

Lastly, recalling the performance of PDE10i from Fig. 1, wherein the drug only recovered 27% of the total distance of the empirically observed WT to HD distance metric constructed from three passive membrane properties, we note that the majority of our virtual drugs performed better in rescuing the WT phenotypes (Fig. 7D).

## Discussion

### Motivation

In contrast to traditional target-based drug discovery strategies, which identify and validate specific molecular targets, phenotypic drug discovery strategies focus on first collecting physiologically relevant end points and then probing for underlying molecular targets in an agnostic manner, thus diminishing target validation risk (49). With more first-in-class drug discoveries being made from such phenotypic drug discovery strategies, a huge opportunity lurks in developing screening tools that enable decision making on the viability of compounds along their well-known multidimenional paths from disease and risk to health and safety, and ultimately for fostering successes in the development of therapies for CNS disorders among the current drug development pipeline.

### HD Phenotypes

Our study began with the assumption that the underlying variability of neuronal electrophysiological phenotypes in HD arises from differential modulation of ion channel conductance densities across principal neurons in the tissue most affected by HD, the striatum. Evidence also points to other sources of variability associated with MSN dysfunction beyond somatic ionic conductances, such as morphological alterations of dendritic topology (50) modulation of extracellular K^+^ levels contributed by K^+^ channel dysfunction among astrocytes (51), TrkB signaling pathway modification of K^+^ channel interacting protein KChIP with Kv4.2 channel subunits (38), alterations to synaptic and receptor function (52) as well as other theoretical and modeling studies that corroborated experiments showing disruptions to the balance between excitatory and inhibitory inputs within striatal networks (53). Kinetic parameters of ion channel models’ sub-threshold voltage gating mechanisms were not varied in this study, though the coregulation of channel kinetics provide general mechanisms for regulating neuronal excitability (54). Prior immunohistochemical studies in MSNs from transgenic mice attributed the reduction in inward and outward K^+^ conductances, as well as in expression of specific K^+^ channel subunits, to the contribution of altered active and passive membrane properties (26) Despite these potentially confounding variables, by varying only maximum conductance parameters our study produced PoMs that closely matched experimental data and provided key methods for measuring drug efficacy and designing multi-target virtual drugs. A summary of our findings includes:

i. Our model-based observations show how reduced K^+^ channel expression can sufficiently alter neuronal excitability (Figs. 3-7) to explain observations that voltage gated K^+^ channels influence membrane depolarization and control striatal neuron firing (55),(10).
ii. In our models, fast and slow inactivating A-Type K^+^ conductances are greatly modified, which contributed to the difference in somatic excitability among WT and HD PoMs’ phenotypes (Figs. 4-6). However, prior evidence of fast inactivating K^+^ channel Kv4.2 dysfunction was localized to distal dendrites of the indirect pathway MSNs (57). Contrary to our findings, in these studies Kv4 current was elevated to make the dendrites hypoexcitable, which occurred alongside decreased cortical drive, through impaired TrkB signaling. We propose that somatic excitability may be an additional homeostatic response to decreased efficacy of cortical drive. This hypothesis requires further experimentation to test, wherein the non-homogenous distribution profile of these ion channels is probed to examine if indeed Kv4.2 are differentially modulated in HD between soma and distal dendrites.
iii. Our parametric analysis of the PoMs revealed a decrease in slowly inactivating K^+^ conductance, which substantiates prior evidence revealing downregulation of Kv2.1 channel expression (26). Several types of K^+^ channel subunits identified, including Kv1.4, Kv4.2, Kv2.1 (58) give rise to slowly inactivating currents. Most likely, the strongest reduction in A-type among the HD PoM, fast and slow inactivating K^+^ and persistent K^+^ conductances (Fig. 3), could be attributed to their crucial role in delaying threshold excitation in the model (TFS feature used in this study) in response to injected current, which captures a key characteristic of MSNs. We imposed a stricter constraint on WT models to reproduce this feature with TFS>100ms, which may have caused these currents to be strongly upregulated in the WT PoM. Furthermore, channel engagement at the different phases of membrane excitation during subthreshold membrane depolarization contributes differentially to this characteristic delayed excitation (59)(42).
iv. In our models, a transient Na^+^ conductance required upregulation in order to rescue the HD phenotype. Though there is no direct evidence of reduced expression of the Nav1.2 channel in HD to our knowledge, evidence that sodium channel β4 subunit downregulation in HD transgenic animals may underlie neuritic degeneration does exist (39) In fact, among cultured cerebellar granule cells, knockdown of Navβ4(Scn4b) revealed the loss of resurgent current, reduced persistent current, and a downward shift in half-inactivation voltage of transient current, thus altering firing patterns. Though, our findings do not establish a direct link to the upstream genetic pathways, reduced Na^+^ currents were not sufficient alone in rescuing cell excitability of HD phenotypes (Fig. 4C).
v. Our models reveal that KRP and NaS were strongly correlated in both WT and HD phenotypes in order to maintain targeted firing properties (Fig. 3B). NaS is a TTX insensitive Na^+^ current and is a known target for modulation of neuronal firing properties (41), while persistent components of the total K^+^ current were pharmacologically heterogeneous, being available over a broad range of membrane potentials (42). It is not surprising that these two currents were strongly implicated in maintaining firing rates across the PoMs.
vi. Surprisingly, the KIR conductances in our HD models were upregulated, while prior evidence attributed hyperexcitability of D2 MSNs from HD model animals to reduced KIR currents. This finding, we believe, was due to the limitations of the protocols we employed for modeling hyperpolarizing current injections, which may have been insufficient to engage KIR conductance modulation of membrane properties to the same extent as experimental protocols (60,61).
vii. In our models, we observed no significant differences between the WT and HD phenotypes with respect to specific leak components. We note that replacing the original nonspecific linear leak model with a set of specific nonlinear leak components (see Methods) was appropriate for overcoming a major drawback of Mahon et al., model (28), i.e., the use of different reversal potentials (E_K_ or E_Na_) for each model channel conductance. Without this modification, biophysical interpretation of the model would be impossible. Secondly, not separating the leak components would have biased the optimization of membrane depolarization towards modulating this leak component, creating degenerate solutions and making it difficult to interpret the influence of specific ionic conductances. Prior modeling work and concomitant patch clamp recordings in Purkinje neurons revealed that GHK based leak models follows a nonlinear I-V relationship and are better at predicting the nonlinear voltage responses to current injections (62).

### Degeneracy

A caveat while interpreting some of our observations from the models is the existence of degenerate solutions of complex systems (63). This has been demonstrated in prior experimental and theoretical studies in neurons, where multiple solutions of ion channel conductances produce similar neuronal excitability phenotypes (64,65). If one channel is deleted, the excitability can still be maintained by other compensatory mechanisms. It is therefore likely that using a different biophysical model of the MSN with more dendritic complexity and ionic conductances contingent upon neuronal topology, such as the Wolf model (67), would reveal alternate virtual drugs and create a more complete reverse screen for recovery. These predictions require further validation in models constrained by detailed measurements, but do not undermine the applicability of our methodology to drug design given appropriate models and appropriate preclinical data sets.

### DARPP-32 and dichotomous MSN types

An important consideration not included in this modeling study is a distinction between different classes of MSNs. Previous studies attributed intrinsic excitability differences to dichotomous D1 and D2 type dopamine receptors (D1Rs and D2Rs) that modulate different classes of MSNs and affect how they process synaptic input integration (67). While dopamine’s regulation of Kir channels in D1R expressing MSNs enhances resonant frequency and reduces resonant impedance, its effect on D2R expressing MSNs is opposite. Each receptor type engages different subcellular biochemical cascades, with D1Rs acting via cAMP-PKA signaling, and D2Rs modulating KIR channels through PLC-PKC signaling (61) These important considerations will drive future studies in which D1-type and D2-type MSN phenotypes are represented in PoMs.

Downstream of these cAMP and cGMP activating pathways is the phosphoprotein DARPP-32, which is highly enriched in MSNs and plays a central role in neurotransmission and in regulating excitability (68,69). Drugs of abuse such as cocaine and amphetamine increase DARPP-32 phosphorylation in the striatum (70,71) and have been used to study the neurophysiology of addiction. The molecule also is central to certain studies targeting neuropsychiatric and neurodegenerative disorders (72). Seminal work by Paul Greengard provided understanding of how various physiological and behavioral effects of antipsychotic drugs and drugs of abuse are greatly reduced in mice with a targeted deletion of the DARPP-32 gene (25). Prior patch-clamp studies quantified the decrease of voltage-gated sodium currents (73) by direct injection of phosphorylated DARPP-32. With such a crucial role for DARPP-32 in regulating neuronal excitability, future work will involve investigating our results in the context of ion channel target modulation by DARPP-32 phosphorylation and its modulation by different drugs. By overlapping a drug’s mechanism of action from high content screen measures and electrophysiological recordings (24,74) with a systems biology model capturing the D1 and D2 receptor modulation of DARPP-32 substrate via PKA and PLC signaling pathways (75), the mapping may provide a way to quantify the effect of DARPP-32 phosphorylation ion channel modulation of membrane activity, and this is therefore a promising direction for future investigation.

### Virtual Drug Translation

*Ex vivo* evaluations supported the notion that dysregulation of cyclic nucleotide signaling can be restored with PDE10i (76). In the symptomatic HD mouse models Q175 and R6/2, cyclic nucleotide dysregulation contributes to neurophysiological dysfunction. PDE10i reversed hyperexcitability of MSNs both *in vivo* and *in vitro*, while also elevating cAMP levels. One particular compound, PF-02545920, a select PDE10A inhibitor, (9) failed to meet the clinical end point of alleviating motor symptoms despite its success at addressing preclinical HD deficits (9). The current study aimed to provide a quantitative measure of how well the drug addressed the recovery of *in vitro* electrophysiological properties in the multidimensional measurement space of MSN phenotypes, as independent statistical comparisons across three features may not have been sufficient to assess the preclinical performance of the drug. We introduced the multidimensional metric, Euclidean distance measure in 3D, and we aimed to recover a heterogeneous population according to divergence measured in an extended metric space (Fig. 1). We set stringent criteria for recovery of the HD PoM’s phenotype into a narrow heterogenous WT region of feature space. Using virtual drugs, we modulated the underlying model parameters and thus introduced novel standards of defining effective phenotypic recovery for both virtual and real drugs.

Though by no means trivial, identifying ideal drug compounds or drug combinations, which target ion channels such as found in our multidimensional targets, is feasible given that screening methods in medicinal chemistry are capable of testing precise modulation profiles. We have identified and presented here methods to measure these profiles against ideal virtual drug profiles (Figs. 7C, D). We also propose searching known drug interaction databases for potential novel compounds that target proteins whose modulation is correlated with elements of our perturbation vectors (i.e., ionic conductance modulation ratios). Finding combinations determined by simple vector arithmetic, a match to the perturbation profile of our best virtual drugs (Figs. 7C, D) becomes possible. The first step for accomplishing this is to map required changes for each ionic conductance (i.e., elements of the perturbation vector) to an equivalent drug dose via the IC_50_/EC_50_ responses of drugs targeting each channel current. In cardiac electrophysiology simulations for pro-arrhythmic safety testing, drug-block within models is achieved by scaling the maximum conductance of an affected ion channel using IC_50_/EC_50_ scaling factors (77). If these values were well catalogued for neuronal ion channels, our proposed virtual drugs could potentially be transcribed into sets of multi-compound therapeutics that are predicted to maximize phenotypic recovery.

### Multi-scale extensions

Unlike in cardiac electrophysiology, where ion channel dysfunction sufficiently explains phenotypic variability (78), in neuronal electrophysiology evidence points to sources of phenotypic variability arising from dendritic topologies, network topologies, and subcellular mechanisms. Our neuronal optimization framework doesn’t restrict construction of such topologically constrained large-scale brain architectures, as the underlying neural tissue simulator is built to handle simulations of high order components, for example a million neurons and a billion synapses (79). A preceding study from our group, using the same optimization methods, employed a model that included dendrites and novel dimensionality reduction techniques to identify metaparameter controllers of sub-threshold oscillations and spontaneous firing in dopamine neurons from a 36-dimensional parametric space (36). Somatic conductance changes determined from single neuron analyses can be easily extended into network architectures to design studies targeting network activity as a phenotype (53) which will be the focus of future studies. With highly parameterized models, careful consideration should be made towards uncertainty quantification of the model parameters in estimating qualitative output behaviors (80).

## Conclusions

In conclusion, we have developed a reverse phenotypic screening method to identify the ‘ideal’ virtual perturbation underlying ion channel targets to rescue a heterogenous population of HD models’ neuronal excitability phenotype to WT. Our approach is based on quantitative systems pharmacology principles, and combines mechanistic simulations, generation of populations of models, and statistical screening approaches for early validation of virtual drug candidates. First, by building databases of neuron models representing both healthy and disease phenotypes we were able to provide mechanistic insights into how the ionic conductance parameters influenced the varied electrophysiological properties that gave rise to distinct clusters of phenotypes. Second, we described several approaches using statistical and machine learning methods to perform reverse phenotypic screening and design single target and multiple target perturbations for rescuing a heterogenous disease population to the healthy phenotype. We also introduced several metrics to compare the performance of the virtual drug perturbations with a known pharmacotherapy using PDE10i in HD. These screening approaches provide novel tools for drug design when complemented with directed *in vitro* electrophysiological studies for preclinical validation of drug targets.

With increasing cost of clinical trial failure in the domain of nervous system disorders and a paradigm shift in drug discovery towards automated high-throughput screening platforms, our approaches may become important for the evaluation of drug action on underlying physiology. As multi-parametric subcellular responses are regularly measured using multiple data streams, from gene expression studies to human-derived iPSC electrophysiological recordings, we anticipate that these concepts will become even more relevant as drug development moves towards provisioning of patient-specific precision therapies (77).

## Study Highlights

### WHAT IS THE CURRENT KNOWLEDGE ON THE TOPIC?

Pre-clinically, potential drug candidates are screened for alleviation of certain *in vitro* electrophysiological features. Population based modeling is widely used to represent intrinsic variability of these features. However, the means by which these drugs can address recovery in terms of divergence of a heterogenous population is neglected and unexplored.

### WHAT QUESTION DID THIS STUDY ADDRESS?

Can computational models of striatal medium spiny neurons, which capture statistical relations between electrophysiological features of Wild-type and Huntington’s Disease phenotypes, and their respective model parameter sets provide a direction of ion channel target modulation for phenotypic recovery across a heterogenous population?

### WHAT THIS STUDY ADDS TO OUR KNOWLEDGE

The methods presented here help identify ‘one’ virtual drug or a profile of target modulations that transform disease phenotypes to healthy phenotypes in terms of both heterogeneity and divergence of the model population.

### HOW THIS MIGHT CHANGE CLINICAL PHARMACOLOGY AND THERAPEUTICS

Complementing pre-clinical experiments with these computational methods can be used to screen potential therapeutic targets. Additionally, also identifying synergistic drugs that align with the target modulation profile to alleviate disease phenotypes in terms of both heterogeneity and divergence at the population level becomes possible.

## Acknowledgments

All authors are employees of IBM Research. Authors would like to acknowledge CHDI Foundation.

## Author contributions

SA, JK wrote the manuscript. SA, TR, JK conceived the design. SA, THT developed the MSN model and TR, THT designed the optimization framework. SA performed the calculations. SA, TR, JP analyzed the data. JK was in charge of overall direction and planning. All authors reviewed the manuscript.

## Code and Data Availability

https://github.com/sallam-usc/Study_Data.git

## Notes

### Competing Interest Statement

The authors have declared no competing interest.

