## Supplemental Methods 1.1 for "Neuronal Population Models Reveal Specific Linear Conductance Controllers Sufficient to Rescue Preclinical Disease Phenotypes"

### Supplementary Methods

#### 1.1 Neuron model

The model comprised eight conductance parameters, the transient sodium current (gNat), persistent sodium current (gNap ), the slowly inactivating sodium current (gNas), the delayed rectifier potassium current (gKDR), the inward rectifying potassium current (gKIR), the persistent potassium current (gKRP), the fast inactivating A-current (gKAf), the slow inactivating A-current (gKAs). The model also included three specific leak currents, for potassium, sodium and chloride, each based on the Goldman-Hodgkin-Katz (GHK) equation:

$$\phi_s = P_s z_s^2 \frac{V_m F^2}{RT} \frac{[S_i]-[S_o]\exp(-z_s V_m F/RT)}{1-\exp(-z_s V_m F/RT)},$$

where $\phi_S$ is the current density flux (amperes per unit area) of ion S, $P_s$ is the permeability of ion S, $[S_i]$ is the intracellular concentration of ion S, $[S]_o$ is the extracellular concentration of ion S, and $V_m$ is the membrane potential. Reversal potentials for each ion species are then calculated once for all currents using the Nernst equation:

$$E_s = \frac{RT}{zF} \ln\left(\frac{S_o}{S_i}\right),$$

where R is the universal gas constant 8.314 J·K$^{-1}$·mol$^{-1}$, T is the temperature 295 K, z is the valence of the ionic species, and F is the Faraday's constant 96,485 C·mol$^{-1}$. We set external and internal ion concentrations to those used for the experiments reported herein in preceding sections. The calculation of MSN membrane potential in our revised model then follows:

$$C\frac{dV}{dt} = -[(V - E_{Na}) * \bar{g}_{NaT} + (V - E_K) * \bar{g}_{KDR} + (V - E_K) * \bar{g}_{KIR} + (V - E_K) * \bar{g}_{KRP} + (V - E_K) * \bar{g}_{KAf} + (V - E_K) * \bar{g}_{KAs} + (V - E_{Na}) * \bar{g}_{NaS} + (V - E_{Na}) * \bar{g}_{NaP} + I_{Cl,leak} + I_{Na,leak} + I_{K,leak} + I_{Inj}),$$

where V is the membrane potential, each $g$ is the conductance of an ion channel current (noted by subscript). Ion channel currents followed $I = \bar{g}m^k h(V - E)$. The activation m and optional inactivation h gating variables were as reported in Mahon et al. 2000 followed:

$$\frac{dp}{dt} = \alpha_p(1 - p) - \beta_p p, \qquad p \in (m, h, n)$$

For channel Nat and KDR, the gating parameters were as follows:

$$\alpha_m = 0.1(V-28)/\exp((0.1(V-28)) - 1), \ \beta m = 4.0\exp(V-53/18),$$

$$\alpha h = 0.07\exp(0.05(V-51)), \ \beta h = 1/(\exp 0.1(V-21) + 1)$$

$$\alpha_n = 0.01(V-27)/\exp((0.01(V-27)) - 1), \ \beta n = 0.125\exp(V-37/80)$$

The inactivation for currents followed:

$$\tau(V) = \tau_0 \Big/ \left(exp^{-(V-V\tau/k\tau)} + exp^{(V-V\tau/k\tau)}\right),$$

except for $I_{As}$, where $\tau_{hAs}(V) = 1790 + 2930\exp(-V+38.2/28^2)((V+38.2)/28)$, and $I_{KRP}$, where $\tau_{KRP}(V) = 3*\tau_{hAs}(V)$. $I_{leak}$ for each specific leak for is derived by multiplying current density flux from equation 1 by the compartment surface area (assumed to be 1 $\mu$m2). $I_{Inj}$ represents injected current.

For numerical integration, we used a time step of 0.01 ms. All datasets were archival at the time they were shared with researchers from IBM, and no new experiments were suggested, designed, or performed based on these analyses.

**1.2 Software**

All simulations were performed using the IBM Neural Tissue Simulator (NTS) on IBM's Cloud. NTS executes simulations based on model descriptions (written in the Model Description Language) and resource allocation scripts (written in the Graph Specification Language). The software is experimental, and readers are therefore encouraged to contact the authors if interested in using the tool. The MSN model is available to run simulations with the parameters sets used in the study are available upon request. The parameter sets to reproduce the wild-type and huntington's disease electrophysiological properties are available to run the simulations.

**1.3 Population modeling**

The optimization employed the non-dominated sorting (NS) differential evolution (DE) algorithm (NSDE) (81,82) previously used to search parameter space of compartmental neuron models in Rumbel et al. (36,83). To run the algorithm, we used a modified version of the BluePyOpt (29). Python framework for single neuron optimization. Trial-and-error was used to assess optimization metaparameters such as population size and number of generations. A single optimization of ~500 generations of a population of ~100 models took approximately ~12 hours of computing time on an X86_64 Intel architecture (2 Ghz, 64 bit, 56 cores, 128 Gb of RAM).

Neuron model error score for each target feature was calculated by extracting feature measures and subtracting them from the exact target values based on empirical measures and dividing the absolute value of this quantity by a deviation variable based on variability of the experimental measures. Dominance ranking according to the NS algorithm was used as the first criterion for model selection, and total error was used to sort models within dominance ranks.

#### 1.4 Parameter search

**Features and stimulation protocols** included those to extract membrane resistance from voltage traces elicited by a 5pA depolarizing stimulus for a duration of 200ms. Rheobase was determined using a ramp protocol with a delay of 500 ms and current gradually increased from 0 to 1000pA over 1000msec, effectively with a slope of 1pA/ms.

A large proportion of models enter into depolarization block. Although certain features are calculable, the results are not deemed accurate, and under further depolarization, not sustainable. Hence additional feature constraints such as interspike interval coefficient of variation (ISI_CV) and firing rates were added. We made multiple checks so that the firing rate is also captured within the time window of the last 1000 ms.

To capture firing rate feature values (spikes/s), and in accordance with data from Planert et al., (21) three separate protocols were used: firing rates at current injection equal to Rheobase, at Rheobase+50pA (FR50), and at Rheobase+100pA (FR100). In addition, ISI_CV within an interstimulus interval, after hyperpolarizing potential (AHP), action potential height (AP_height), time to first spike, or spike latency when elicited by Rheobase+50pA and Rheobase+100pA were each targeted as features constrained by the optimization algorithm. Two separate runs of optimizations were performed for WT and HD categories.

It was difficult to find PoMs spanning the complete feature ranges uniformly. We stopped the optimization when we achieved >1200 zero error models. The WT and HD firing rates were targeted to a mean value of 6 spikes/s and a target deviation of 4 spikes/s. In this way, the optimization algorithm would accept a model as a 'good' model if it had a firing rate between 2 - 10 spikes/s.

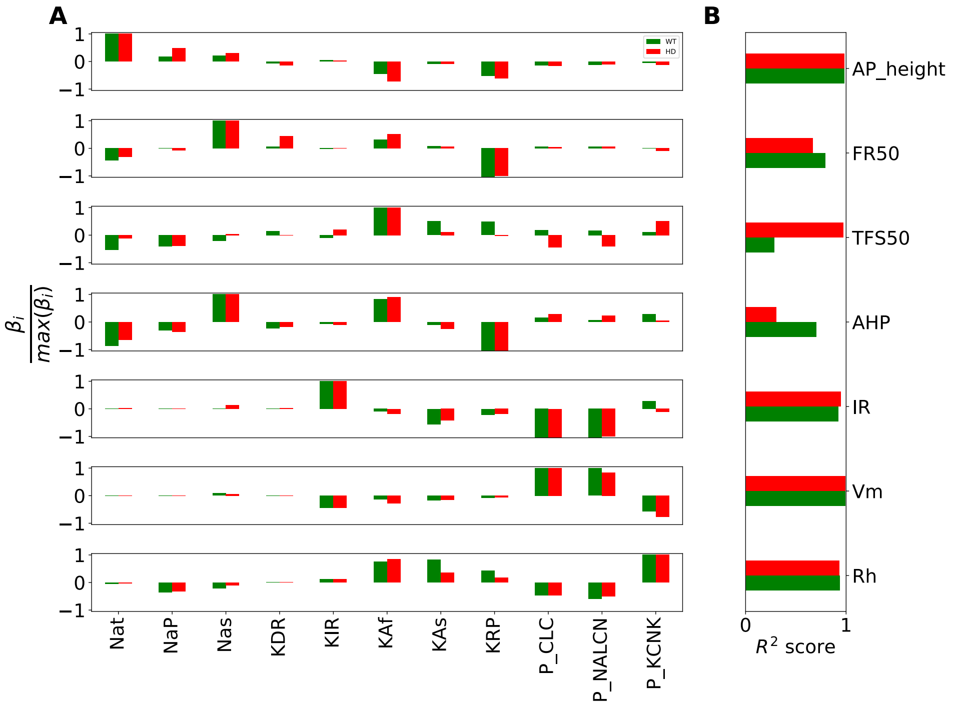


Figure S1. **Linear regression analysis of WT and HD population of models.** A) MSN electrophysiological features are modulated by the underlying 11 ionic conductances and a subset of these parameters are strong modulators of the features. Side-by-side comparisons of parameter coefficients from WT (green) and HD (red) phenotypes. B) Goodness of fit measure of the linear model for each feature.
